# Striking differences in patterns of germline mutation between mice and humans

**DOI:** 10.1101/082297

**Authors:** Sarah J. Lindsay, Raheleh Rahbari, Joanna Kaplanis, Thomas Keane, Matthew E. Hurles

**Affiliations:** Wellcome Sanger Institute, Hinxton, Cambridge, CB10 1SA, UK

## Abstract

Recent whole genome sequencing (WGS) studies have estimated that the human germline mutation rate per basepair per generation (∼1.2−10^−8^) ^1,2^ is substantially higher than in mice (3.5-5.4−10^−9^) ^3,4^, which has been attributed to more efficient purifying selection due to larger effective population sizes in mice compared to humans.^5,6,7^. In humans, most germline mutations are paternal in origin and the numbers of mutations per offspring increase markedly with paternal age ^2,8,9^ and more weakly with maternal age ^10^. Germline mutations can arise at any stage of the cellular lineage from zygote to gamete, resulting in mutations being represented in different proportion and types of cells, with the earliest embryonic mutations being mosaic in both somatic and germline cells. Here we use WGS of multi-sibling mouse and human pedigrees to show striking differences in germline mutation rate and spectra between the two species, including a dramatic reduction in mutation rate in human spermatogonial stem cell (SSC) divisions, which we hypothesise was driven by selection. The differences we observed between mice and humans result from both biological differences within the same stage of embryogenesis or gametogenesis and species-specific differences in cellular genealogies of the germline.

## Main text

To characterise mutation rates, timing and spectra in the murine germline we analysed the patterns of *de novo* mutation (DNM) sharing among offspring and parental tissues in six large mouse pedigrees (Extended Data Figure 1), using a combination of WGS, deep targeted sequencing and an analytical workflow described previously^9^. We then compared these murine patterns with equivalent, previously published, human data on three multi-sibling families^9^. After WGS, discovery and validation of candidate DNMs in 5 or 10 offspring from each mouse pedigree, we validated 753 unique autosomal DNMs (746 SNVs, 7 MNVs (multinucleotide variants), with a range of 8-36 *de novo* SNVs per offspring (Extended Data Table 1). The validated mutations were genotyped in other offspring from each pedigree (Figure 1). We determined that 2.7-fold more unique DNMs were of paternal (N=152) than maternal (N=55) origin, similar to previous studies^3,4^. Mice and humans have more similar paternal biases in mutations than might be expected (2.7:1 and 3.9:1 respectively^2,8,9^), given the five-fold difference in the ratios of genome replications in the paternal and maternal germlines between mice (2.5:1) and humans (13:1)^11^

**Table 1:** Autosomal SNV germline mutation rates per generation, per year and per cell division in humans and mice. 95% confidence intervals calculated assuming Poisson variance around the mean number of mutations (Methods). Rate per cell division are calculated assuming that mouse and human generation times are 9 months and 30 years respectively and the numbers of cell divisions delineated in reference^13^. Estimates of mouse mutation rates per year are highly dependent on estimates of average generation time.

|  | Human | Mouse |
| --- | --- | --- |
| Mutations per genome per generation | ~71 | ~20 |
| Mutation rate per genome per generation | $1.22 \times 10^{-8}$ [$1.14 \times 10^{-8}$ - $1.31 \times 10^{-8}$] | $0.39 \times 10^{-8}$ [$0.37 \times 10^{-8}$ - $0.42 \times 10^{-8}$] |
| Mutation rate per year | $4.08 \times 10^{-10}$ [$3.80 \times 10^{-10}$ - $4.35 \times 10^{-10}$] | $53 \times 10^{-10}$ [$49 \times 10^{-10}$ - $56 \times 10^{-10}$] |
| Average mutation rate per cell division | $5.67 \times 10^{-11}$ [$5.28 \times 10^{-11}$ - $6.05 \times 10^{-11}$] | $9.07 \times 10^{-11}$ [$8.41 \times 10^{-11}$ - $9.69 \times 10^{-11}$] |

**Figure 1:**
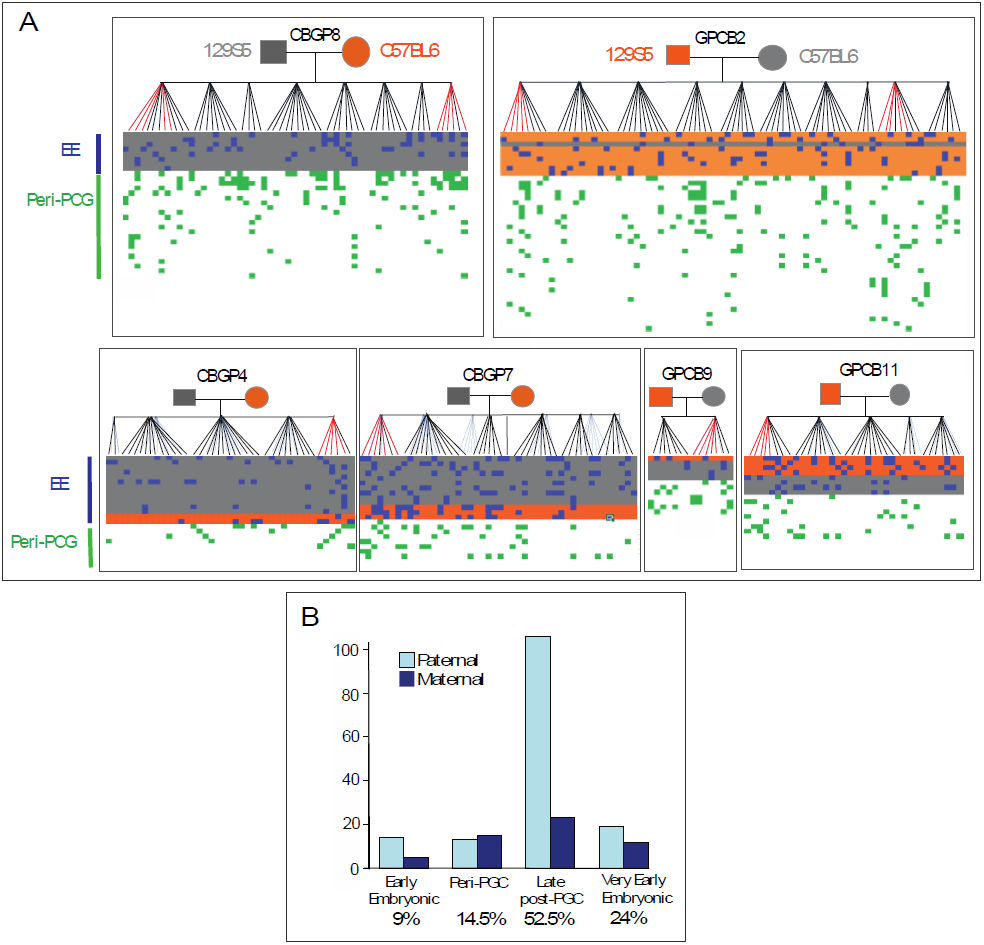
A) Validated *de novo* mutations in six mouse pedigrees. Offspring are shown vertically, and DNMs are shown horizontally, coloured according to temporal strata. Early embryonic DNMs are shaded according to the parental origin (grey/orange). Late post-PGC and VEE mutations are only observed in one individual and are listed in Extended Data Table 1. **B)** The number of DNMs assigned to the paternal or maternal haplotype (Methods) in each temporal strata, with the total contribution of mutations in each strata to the overall rate shown at the bottom of the graph.

In the two largest mouse pedigrees, we observed 18% (70/388) of unique DNMs were shared among 2-19 siblings, strongly implying a single ancestral mutation. The fraction of mouse DNMs that are shared between two siblings is significantly higher (p=4.0−10^−7^, proportions test) in mice (3.1%) than has been reported in humans (1.2%)^9^, suggesting that a higher proportion of DNMs in mice derive from early mutations in the parental germline leading to mosaicism in the parental germline (Extended Data Table 2).

For both species, using the patterns of mutation sharing across the parental and offspring tissues, we classified DNMs into four temporal strata of the germline (Extended Data Figure 2A, Extended Data Figure 2B). We refer to these four strata as very early embryonic (VEE), early embryonic (EE), peri-primordial germ cell specification (peri-PGC) and late post-primordial germ cell specification (late post-PGC). VEE mutations likely arose within the earliest post-zygotic cell divisions contributing to the developing embryo, EE mutations arose during later embryonic cell divisions, prior to PGC specification (after ∼10 cell divisions), peri-PGC mutations arose around the time of PGC specification, and late post-PGC mutations arose during PGC proliferation and gametogenesis. Collectively, these four temporal strata represent a full generational span (Extended Data Figure 2B), albeit with the VEE mutations being quantified here in offspring. VEE mutations accounted for 23.9% (194/811) of all observed DNMs in mice but only 4% in humans (28/719) (Figure 1). The Variant Allele Fraction (VAF) for the observed VEE mutations in mice and humans were consistent with the vast majority occurring in the first cell division that contributes to the embryo and were highly concordant between different tissues. The number of VEE mutations per offspring varied considerably more than expected under a Poisson distribution (p=0.0019), suggesting this stage is more mutagenic for some zygotes than others.

**Figure 2:**
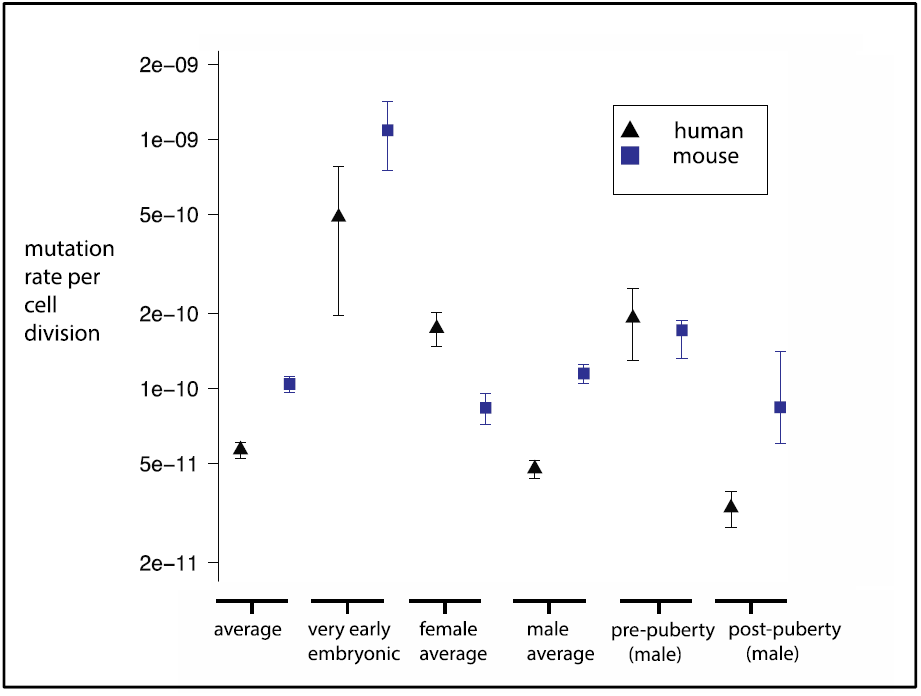
Estimation of mutation rates per haploid base per cell division. Mean per-generation mutation rates for SNVs were calculated for mice and from humans using all the data from this study and Rahbari et al^9^ and Kong et al^2^. The 95% confidence intervals were calculated assuming that DNMs are Poisson distributed, except for the pre and post-puberty stages which were derived from the linear models fitted for the paternal age effects, and VEE mutations which assume quasi-Poisson distribution to allow for over-dispersion. See Methods for further information. The accuracy of the estimates shown here (aside from VEE mutations) rely upon the proposed cellular demographies by Drost and Lee^11^.

The 55 EE mutations we detected in mice were present at similar levels across three parental somatic tissues (1.6-19%), and we observed a significant, but modest correlation between levels of somatic and germline mosaicism (Pearson correlation coefficient of 0.40, p=0.0025, Extended Data Figure 3, Extended Data Table 3). We observed a significant paternal bias among the EE mutations identified in mice (41 paternal, 14 maternal, p=0.0004, two-sided binomial test), but not in humans (9 paternal, 16 maternal, p=0.23, two-sided binomial test), and this difference between species is significant (p=0.002, Fishers Exact Test). We propose two possible biological explanations for this paternal bias in mice: (i) an elevated paternal mutation rate per cell division or (ii) a later paternal split between soma and germline (i.e. more shared cell divisions). Further work is required to distinguish between these two scenarios, although the observation of early sex dimorphism in pre-implantation murine and bovine embryos^12,13^ may well be relevant. The high proportion (average 30%) of DNMs in mice that appear to occur during early embryonic development (VEE+EE mutations), is in accordance with high estimates of germline mosaicism from phenotypic studies^14,15,16^

**Figure 3:**
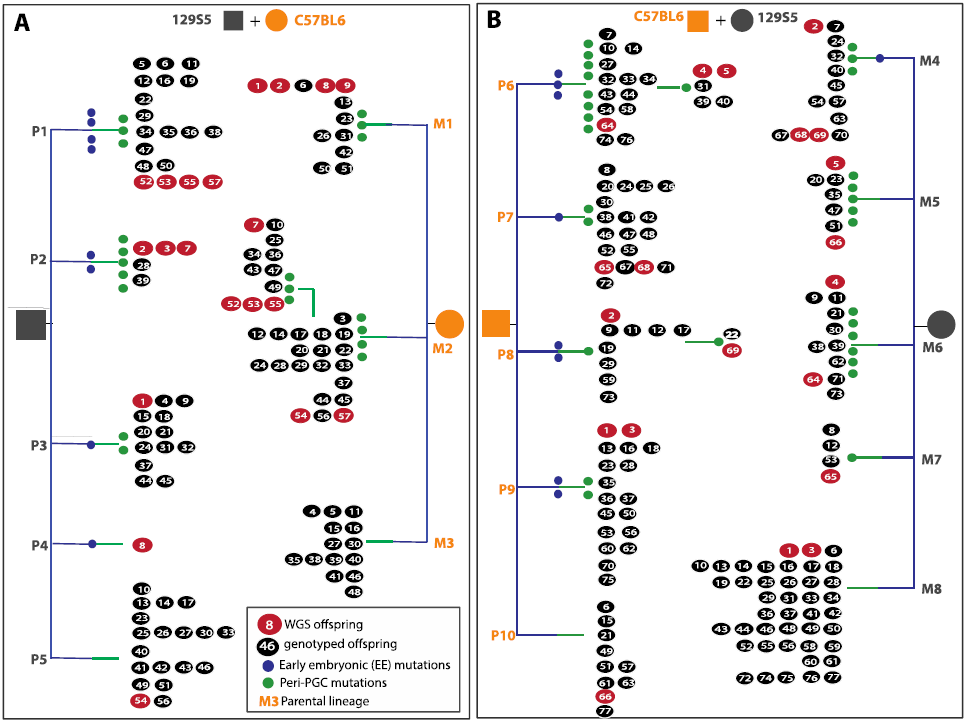
Genealogies of maternal and paternal cell lineages delineated by early embryonic and peri-PGC mutations.

We identified 54 peri-PGC DNMs (shared among siblings but not detectable in parental somatic tissues) in the two largest mouse pedigrees, but with no sex bias in parental origin, where it could be determined (25 maternal, 25 paternal), suggesting that the sex differences identified for EE mutations are confined to a small developmental window.

For both species, we estimated germline autosomal mutation rates per generation, per year and per cell division (Table 1), taking into account the partial contribution of mosaic (VEE) mutations to the germline (Methods). We estimated the mutation rate per generation in mice to be approximately one third that in humans, whereas the annual mutation rate was 13 times higher and the per cell division mutation rate was more than one and a half times higher. The 13-fold difference in annual mutation rates between extant mouse and human is substantially greater than the approximately two-fold greater accumulation of mutations on the mouse lineage reported since the split from the human-mouse common ancestor ∼75 million years ago^17^. However, comparing the annual mutation rates inferred from the human-chimp and mouse-rat sequence divergence (1.3% and 17%) and age of their most recent common ancestors (∼6MYA and ∼12MYA) suggests a seven-fold difference, only a factor of two different from our estimate ^18^. More accurate estimates of average generation times may help to resolve this discordance.

We observed significant differences (p=1.27−10^−9^, chi-squared test) in the mutational spectra in mice and humans^9^. These differences are predominantly characterised by an increased proportion of T>A mutations, and a decreased proportion of T>C mutations in mice (Extended Data Figure 4A(i)). Mice exhibited a stronger mutational bias towards AT bases than humans (69% vs 59% of all such mutations), in accordance with previous studies that have suggested that GC content is decreasing more markedly in mouse genomes^19^. We observed significant differences (p=2.2×10^−7^, Chi-squared test) in the murine mutation spectra apparent in earlier (VEE + EE) and later (peri-PGC and late post-PGC) stages of the germline (Extended Data Figure 4A(ii)), suggesting differences in mutation processes between embryonic development and later gametogenesis.

In mice, we observed a significant (p=0.004) increase in the average number of pre-zygotic mutations (discounting VEE mutations) per offspring with increasing parental age. As in humans, the parental age effect in mice is likely to be predominantly paternally driven, as pre-zygotic mutations exhibit the greatest paternal bias (4.3:1 compared to 2.7:1 overall). The rate of increasing mutations with parental age observed in mice is approximately five-fold greater (p= 0.0003) than in humans, which is larger than the two-fold more rapid rate of turnover of SSCs in mice compared to humans, implying a higher mutation rate per SSC division in mice compared to humans^11^.

We calculated mutation rates per cell division at different phases of the germline in humans and mice (Figure 2), by integrating information on the cellular demography of the germline^11^, the paternal age effect, and the numbers of mutations arising in each temporal strata (Methods). These estimates do not include uncertainty in the numbers of cell divisions per generation or generation times. Mutation rates per cell division are highest in the first cell division of embryonic development in both species. High mutation rates at this earliest stage of embryogenesis is supported by comparable studies in cattle.^13^ The most striking difference between the species is the much lower mutation rate in SSC divisions in humans. SSC cell divisions are significantly less mutagenic than all other germline cell divisions in humans, but not in mice. SSC divisions account for >85% of all germline cell divisions in humans but only <40% in mice^11^. We hypothesise that the much greater relative contribution of SSC divisions to the human germline (Extended Data Figure 2C) has led to stronger selection pressures to reduce the mutation rate of SSC divisions in humans than in mice.

Using mutations shared among siblings in the two largest mouse pedigrees, we reconstructed four partial cellular genealogies for each parent (Figure 3). Each parental genealogy is characterised by 2-4 lineages defined by EE and peri-PGC mutations, and a residual group of offspring without shared mutations. An unknown number of PGC founder cells are specified during mouse embryonic development which later generate 40-42 PGCs^20,21,22^. We noted markedly unequal contributions of different parental cellular lineages to gametes (range: 2 to 53%). The M2 lineage accounted for over half of all gametes and yet was defined by four peri-PGC mutations. This suggests that specified PGCs do not contribute equally to the final pool of gametes, although further work is required to determine the relative contribution of selective and stochastic factors.

Parental embryonic lineages reconstructed for each pedigree in the two largest pedigrees (CBGP8/GPCB2) A, as paternal (P) or maternal (M), with each lineage numbered. Mutations delineating the lineages are colour-coded according to their temporal strata are listed in Supplementary Table 1. WGS and genotyped offspring are shown as red and black numbered circles respectively. Offspring are numbered and ordered according to litter; for example, in lineage P2, individuals 2,3 and 7 belong to the same litter, whereas numbers 28 and 39 arose in separate, later, litters. Lineages P5, M3, M8 and P10 represent offspring without shared mutations and may contain several uncharacterised lineages

The pedigree-based study presented here complements differences in mutation processes inferred from comparative genome analyses^23^, but adds sex-specific and temporal granularity. Some of the differences we observed between mice and humans are attributable to the differences in cellular genealogies of the germline, for example, the greater proportion of germline replications occurring during early embryogenesis in mice, leading to greater germline mosaicism and sharing of mutations between siblings. However, other differences represent biological differences within the same stage of embryogenesis or gametogenesis, for example, the striking paternal bias of EE mutations in mice, or the reduced rate of mutation in SSC divisions in humans. Recent work has challenged the paradigm that differences in germline mutation processes between species is dependent on ‘life history’ traits^24,25^. Pedigree-based studies of other species will establish a truly comprehensive molecular and cellular understanding of the evolution of mammalian germline mutation processes.

## Online Methods

### Mice

Ten male and female mice from two inbred lines (CB57BL/6 and 129S5) were obtained from sib-sib inbred lines previously established at the Wellcome Trust Sanger Institute. Twenty breeding pairs were established: ten CB57BL/6 ♂ x ten 129S5 ♀ (GPCB), and ten of the reciprocal cross: 129S5 ♂ x CB57BL/6 ♀ (CBGP). Breeding pairs were introduced for mating at regular intervals, if a pregnancy resulted, the pups were left to wean and then culled at 3-4 weeks of age. Tissue samples of spleen, kidney and tail were taken from pups, and from the parents either when one of them died or became ill, or when no pregnancies resulted after matings over a period of three months. Once all pairs had stopped breeding, the pair from each cross with the longest breeding span and the most offspring were chosen for the initial WGS experiment. Subsequently, four additional pedigrees, two from each of the reciprocal crosses, were chosen undertake sequencing in a supplementary sequencing experiment. At the onset of the experiment, the ages of the selected mice were between 8 and 12 weeks (Extended Data Table 1).

### DNA Sample Preparation

Tissues were stored at −80°C immediately after harvest. DNA was prepared using Qiagen DNeasy kits. Where possible, single DNA aliquots were used for both discovery and validation experiments.

### Sequencing and variant calling

DNA extracted from the spleen of parents and offspring was sequenced using standard protocols and Illumina HiSeq technologies with a read length of 100bp in both parents and ten offspring from the earliest and latest time-matched litters in the two largest mouse pedigrees (GPCB2 and CBGP8). The resultant sequence data was aligned to mouse reference GRCm38. The total mapped coverage after duplicate removal had a mean of 25X and range 22-35X for pedigree CBGP8, and 29X and 22X-40X for pedigree GPCB2. Variants were called using bcftools and samtools and standard settings^26^. Five offspring and the parents from each pedigree in the remaining four pedigrees CBGP7, CBGP4, GPCB11 and GPCB9 were subject to WGS using standard protocols and Illumina X10 technologies, using DNA from the spleen. The total mapped coverage after duplicate removal had a mean of 41X and a range of 41-47X for CBGP4, 38-45X for CBGP7, 39-44X for GPCB11 and 40-42X for GPCB9. Strain and sex specific SNVs were used to confirm the identify of parents.

### *De novo* mutation calling

*De novo* mutations were called on candidate variants supplied by bcftools using *DeNovoGear* version 0.5 using standard settings^27^. *DeNovoGear* called between 7,711 and 11,069 (mean 9,736) candidate *de novo* short indels and SNVs in GPCB2 and CGPB8, and between 6405 and 11071 candidate SNVs and short indels (mean 9478) in GPCB9, GPCB11, CBGP4 and CBGP7. Candidate *de novo* SNVs and indels on the X chromosome exhibited marked strain/sex specific inflation and were discarded.

### Filtering of candidate *de novo* mutations

Candidate *de novo* mutations were filtered to exclude simple sequence repeats and segmental duplications which are sequence contexts highly enriched for false positives. In addition, strain-specific mapping artefacts (low quality areas leading to clustered/low quality SNV/indel candidates were filtered by removing sites that had a high variant allele ratio (>20%) in any offspring in the litter from the reciprocal cross, or parent of the reciprocal cross (>4%). Assuming a Poisson distribution for sequencing depth, sites with a depth greater than the 0.0001 quantile were removed due to the likelihood of mapping errors or low complexity repeats introducing false positives. For the two largest pedigrees (CBGP8 and GPCB2), candidate sites where the *de novo* mutation was present in either parent in greater than 5% of reads and where there were known SNPs in the parental strain were also removed on the grounds that they were likely to be inherited. For the four additional pedigrees (CBGP7, CBGP4, GPCB9 and GPCB11), filtering of candidate DNMs was applied as above, but without any upper threshold for the proportion of variant alleles in either parent at a candidate site. Instead, we calculated a mutation-specific error rate at each candidate site using WGS data from all unrelated individuals. Candidate sites that had mutation-specific error rate of >2% in unrelated individuals were removed. Finally, sites with a low variant allele ratio (<15%) in the candidate offspring were removed. Once these filters were applied, 272, 380, 225, 260, 205, 324, 166, 286, 284, 375 and 211, 174, 180, 346, 135, 101, 160, 143, 191, 300 candidate *de novo* mutations remained for GPCB2 and CBGP8, and 61, 65, 74,1 06, 70, 167, 55, 142, 130, 96, 95, 111, 92, 103, 83, 57, 71, 48, 79, 103 for CGBP7, CBGP4, GPCB11 and GPCB9.

### Experimental validation of *de novo* mutations

We carried out two separate validation experiments, one for pedigrees GPCB2 and CBGP8 and one for CGBP7, CBGP4, GPCB11 and GPCB9. A total of 4,460 unique sites across all 20 offspring in GPCB2 and CBGP8 were put forward for validation by targeted sequencing and 1,809 unique sites in the CGBP7, CBGP4, GPCB11 and GPCB9 pedigrees. (Agilent Sure Select). Twenty-one sites were lost during liftover conversion in the GPCB2 and CBGP8, leading to 4,439 sites put forward for bait design. Bait design for both experiments included 2X tiling, moderate repeat masking, maximum boosting, across 100bp, of sequence flanking the site of interest (extending to 200bp where baits could not be designed on the initial attempt for both experiments. In GPCB2 and CBGP8, 3,253 sites were successfully designed for with high coverage (>50% coverage), 222 with medium coverage (>25% coverage), and 421 with low coverage (<25% coverage). In the four additional pedigrees, 1131 sites were successfully designed for with high coverage (>50% coverage), 11 with medium coverage (>25% coverage), and 0 sites with low coverage (<25% coverage). 564 and 667 sites failed bait design, and are likely enriched for false positives. Target enrichment was performed on DNA prepared from tissue samples from parents and offspring using standard protocols and sequenced using 75bp paired end reads using Illumina Hiseq instruments. We sequenced all candidate DNMs in three tissues from the parents (mean coverage of 400-800X per tissue), two tissues from the WGS offspring (mean coverage of 400X) and a single tissue from all other offspring (mean coverage of 200X). The resultant sequence data were merged by individual and annotated with allele counts at each candidate *de novo* site using an in-house python script. An in-house R script (http://www.Rproject.org) was then used to calculate the likelihood for each candidate variant being a true *de novo* mutation, an inherited variant or a false positive call, based on the allele counts of the parents and offspring at that locus. A proportion of the SNV candidates (all sites put forward for validation for one individual) as well as all of the indel candidates were reviewed manually using Integrative Genomics Viewer (IGV)^28^. Sites that were 5bp or less apart were designated MNVs (Multinucleotide variants) and counted as one mutation.

### Functional Annotation of variants

Functional annotation of validated DNMs was carried out using ANNOVAR^29^. Seventeen DNMs impacted on likely protein function (one nonsense and sixteen missense), however, none were in genes known to have a dominant phenotype in mice, or are associated with somatic driver mutations, and so were assumed to be representative of the underlying mutational processes and not under strong selective pressures (Extended Data Table 4).

### Classification of Temporal strata

#### Identification of Very Early Embryonic mutations in offspring

We combined the allele counts across both sequenced tissues in each WGS offspring, after first checking that the allele ratios were concordant across tissues (Fishers Exact Test). Very Early Embryonic (VEE) mutations were identified using a likelihood-based test on the combined allele counts by comparing the likelihood of the data assuming the mutation was constitutive (binomial p=0.5) or was generated in the first cell division giving rise to the embryo (binomial p=0.25). DNMs with an absolute log likelihood difference of >5 were designated as VEE or constitutive respectively. DNMs with an absolute log likelihood difference of <5 was considered unassigned and accounted for 10% of mutations in human pedigrees, and 4% in mouse pedigrees. VEE mutations were observed mosaic in offspring tissues and absent from parental tissues, with consistent representation in 25-50% of cells across different offspring tissues, reflecting their likely origin within the earliest post-zygotic cell divisions contributing to the developing embryo (Extended Data Figure 5, Extended Data Figure 6). In addition, phasing of VEE mutations with nearby informative heterozygous SNVs, where available, was used to estimate haplotype occupancy (HO), defined as the proportion of the ancestral haplotype carrying the variant allele. Constitutive mutations should have an HO of 100%, whereas VEE mutations will only be seen on a subset of haplotypes. (Extended Data Figure 5). DNMs shared among offspring (which are constitutive by definition) have high HO, whereas putative VEE mutations were enriched for low HO values that correlated with the estimated level of mosaicism. VEE mutations in mice arose at similar rates in both sexes (average per male offspring=5, average per female offspring =6), and approximately equally on paternal and maternal haplotypes (15:9 paternal:maternal) in the largest two pedigrees. Unassigned DNMs were designated as late-post-PGC mutations in downstream analyses.

#### Calculation of Haplotype Occupancy

We looked for heterozygous SNVs with 100bp of validated DNMs in each offspring. We then calculated the phase of the DNM according to the adjacent heterozygous SNV. DNMs that are pre-embryonic should be in complete phase with one or other of the adjacent alleles (Haplotype occupancy =1). DNMs that arose during embryo development will be observed only on a subset of the ancestral haplotype (Haplotype Occupancy <1) (Extended Data Figure 5).

#### Identification and power to detect EE mutations in parents

In order to identify DNMs that could be mosaic in one of the parents, the site-specific error was calculated for each site (% of reads that map to the non-reference variant allele in unrelated individuals from the reciprocal pedigree). This error was then used to calculate the binomial probability of observing ***n*** non-reference reads at the mutated site in each tissue in each individual. The probabilities were corrected for multiple testing, using Bonferroni correction, with a threshold of p<0.05 to identify candidate sites, which were then viewed in IGV^28^. EE mutations were observed constitutively in offspring and mosaic in parental somatic tissues in a lower proportion of cells (2-20%) than VEE mutations, compatible with them arising during later embryonic cell divisions, prior to PGC specification. In addition, the power to detect mosaicism at different levels (0.5%, 1, and 1.5% respectively), in each tissue in each parent was estimated using the sequence depth from the validation data. All EE mutations were observed in all tissues from the mosaic parent). The proportion of DNMs observed as EE mutations observed in mice represented a very similar proportion of all of EE mutations observed in human pedigrees^9^. The correlation between germline and somatic mosaicism was notably stronger among paternal EE mutations than maternal EE mutations, and is suggestive of sex differences in the cellular genealogy of early embryonic development (Extended Data Table 3).

#### Discounting of technical artefacts in assigning parental origin of EE mutations

We considered and discounted a wide variety of possible technical artefacts that might explain the apparent parental sex bias we observe in early embryonic mutations in mice. Firstly, sequencing depth, and thus power to detect somatic mosaicism, was equal between maternal and paternal tissues, and the identity of the WGS samples were checked using strain and sex-specific SNVs. Secondly, where parental origin could be independently determined by phasing with nearby informative sites (N=6), the parental origin was confirmed, thus excluding sample swaps. Thirdly, parental mosaicism in the deep targeted sequencing data was supported by nonzero counts of variant alleles in the WGS data in the corresponding parents at six of the mosaic sites (five paternal, one maternal). Fourth, the same aliquot of DNA was used for WGS and validation by deep targeted sequencing of mutations in parental spleen, lowering the possibility of sample swaps. Lastly, in all cases, parental mosaicism was independently supported by sequencing data from two additional tissues.

#### Identification of peri-PGC mutations

Peri-PGC mutations were observed shared among two or more offspring of the same parents and so are likely parental germline mosaic, but are not detectably mosaic in parental somatic tissues (<1.6% of cells), compatible with them arising around the time of PGC specification and the separation of germline and somatic cell lineages. After specification, PGCs proliferate rapidly, generating thousands of germ cell progenitors in both sexes, so in the absence of strong positive selection, only mutations that occur prior to this proliferation are likely to be observed in multiple siblings in our pedigrees; studies of phenotypic markers of mutation have indicated that spermatogonial stem cells need to be depleted almost to extinction to result in sharing of phenotypes induced by later mutations among offspring14,^15,16^ We confine our analysis of peri-PGC mutations to the two largest pedigrees due to the greater power to detect this class of mutation in the largest pedigrees where a greater number of offspring have been whole genome sequenced. We identified 54 peri-PGC DNMs shared among two or more offspring but not present at detectable levels (>1.6% of cells) in parental somatic tissues of the two largest pedigrees (Figure 1). Peri-PGC mutations arose approximately equally in the paternal and maternal germlines (direct phasing: 6 paternal, 8 maternal; inferred parental origin using co-occurrence: 25 paternal, 25 maternal). We only observed 4 peri-PGC DNMs in the human pedigrees, although the numbers are not comparable between species, due to the disparity in numbers of offspring per pedigree and therefore the power to distinguish this class of DNMs.

#### Identification of late-post-PGC mutations

Late post-PGC mutations were observed constitutively in a single offspring, and represent mutations arising during cell divisions from PGC proliferation onwards.

### Haplotyping of *de novo* mutations in offspring

We used the read-pair algorithm within *DeNovoGear*^27^ to determine the parent of origin of validated DNMs using the whole-genome sequence data. *DeNovoGear* uses information from flanking variants that are not shared between parents to calculate the haplotype on which the mutation arose.

Using this technique, we were able to confidently assign the parental haplotype in 207 of 753 unique validated DNMs.

### Estimating the autosomal SNV mutation rate per generation

Sites that were 5bp or less apart were designated MNVs (Multinucleotide variants) and counted as one mutation. We estimated the number of autosomal DNMs in each mouse offspring by correcting for the proportion of the genome that was interrogated as follows. Bedtools^30^ was used to calculate the proportion of the genome considered in our analysis after removing sites with low- or high-sequence depths for each individual (p_depth_). We then calculated the proportion of sites that were retained after applying our whole-genome filters (simple sequence repeats and segmental duplications) after the depth filters were applied (p_filter_). Last, we used the posterior probability supplied by *DeNovoGear* to calculate what proportion of true DNMs arose at sites that could be validated (p_dnm_). Multisite variants were considered to be a single mutational event. The mutation rate was estimated as follows where 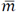 is the average number of mutations observed per offspring.

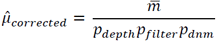

To calculate the mutation rates per generation, per year and per cell division for human and mouse comparisons (reported in Table 1), generation times were assumed to be 30 years and 9 months respectively. The average age for the parents in the human study was 29.8 years^9^, while the parents in the mouse experiment were 24.5 weeks old on average when the offspring were conceived. According to Drost^11^, generation times of 30 years and 9 months would result in 432 cell divisions in the human germline (401 paternal, 31 maternal), and 87 cell divisions in the mouse (62 paternal, 25 maternal). The mutation rate per generation per haploid transmission was then calculated as follows:

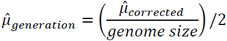

95% confidence intervals were calculated assuming Poisson variance around the mean number of mutations observed. The genome size used was the total number of non-N autosomal sites. The mutation rate per year was then calculated as the mutation rate per generation divided by the average generation time for humans and mice in years.

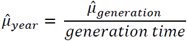

Lastly, the mutation rate per cell division was calculated as the mutation rate per generation divided by the sex-averaged number of cell divisions in a generation^11^, calculated as the sum of the number of cell divisions in the male and female germline divided by two.

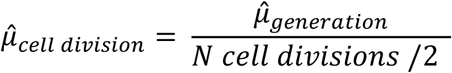

### Estimating contribution of VEE mutations to germline mutations

We identified candidate VEE mutations that occurred in the early embryonic development of the parents in the deep WGS in mouse parents in four smallest pedigrees. We prioritized candidates in these pedigrees as we had greater power to detect parental VEE mutations due to the greater WGS depth compared to the two larger pedigrees. SNVs were called in each pedigree using Platypus^31^. Raw candidates were filtered to remove sites known to be variant in that mouse strain, sites present in Segmental Duplications or Simple Sequence Repeats, and retain only those sites called as heterozygous in a single parent.

Candidate sites were then further filtered to retain only sites where the variant base was observed in fewer than 5% of the total reads in all individuals in unrelated pedigrees. For each remaining site, the mutation-specific error rate was calculated from individuals in unrelated pedigrees. This mutation-specific error rate was used to remove sites with a Poisson probability of >0.02 of being a sequencing error. Sites with a binomial probability of >0.003 of being constitutive were also removed. Finally, to maximise stringency, we removed candidate sites with more than one alternate read in any other parent, fewer than four variant sequence reads in the candidate individual and that the candidate VAF was greater than 35%.

After applying the filters above, 44 candidate sites remained across the four pedigrees. We validated these mutations across three tissues in both parents and a single tissue in all related offspring using a PCR and indexed sequencing approach, described previously^32^. Sites were sequenced (Extended Data Figure 1) to an average of 100,000X coverage across the candidate site of interest in the parents, and >20,000X in the offspring. We assayed all the available offspring from each pedigree, which increased the numbers of offspring in pedigrees GPCB9 and GPCB11 compared to previous analyses (Extended Data Figure 1). The candidate sites were annotated with read counts at the candidate site using an in-house python script. Twenty six sites out of twenty nine with high sequence depth in both parents and offspring were classed as true VEE mutations, on the basis of VAF in the parents being incompatible with either sequencing errors or a constitutive variant. For these 26 sites, the number of offspring to whom the mutation was transmitted was determined (Extended Data Figure 7, Extended Data Table 5).

A linear model was fitted to the outcome of the parental VEE quantification experiment (Methods, Extended Data Figure 7, Extended Data Table 5) to infer the relationship between somatic VAF in parents and proportion of offspring carrying the parental VEE mutation. We observed a significant correlation between the Variant Allele Fraction of the VEE mutation and the proportion of offspring to which the mutation was transmitted (Pearson correlation = 0.71, p=0.00011; Extended Data Figure 7, Extended Data Table 5). For simplicity, we used the point estimate from the linear model, rather than a confidence interval. On average, this calculation reduced the contribution of VEE mutations to the germline mutation rate by 40%.

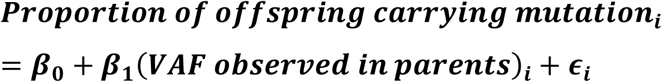

This model was then used to predict the likely germline contribution of VEE mutations observed in offspring given their VAF.

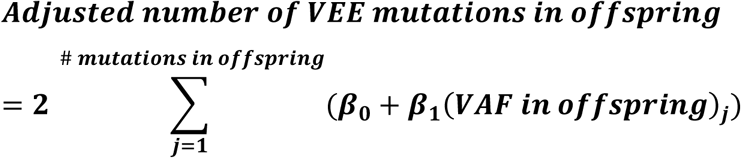

### Analysis of mutation spectra

Mutational spectra were derived directly from the reference and alternative (or ancestral and derived) allele at each variant site. The resulting spectra are composed of the relative frequencies of the six distinguishable point mutations (C:G>T:A, T:A>C:G, C:G>A:T, C:G>G:C, T:A>A:T, T:A>G:T), with the C:G>A:T then split into mutations at a CpG and non-CpG context. Significance of the differences between mutational spectra was assessed by comparing the number of the six mutation types in the two spectra by means of a Chi-squared test (df = 6). The differences between the mutation spectra in mouse and humans cannot be accounted for by the slight difference in genome-wide base composition between human and mouse genomes (GC content of 42% and 41% respectively) as the two most discordant classes of mutation shared the same ancestral base (T) but exhibited opposing directions of change. We observed no significant differences in the mutation spectra between maternally and paternally derived DNMs in mice (p= 0.239, Chi-squared test, Extended Data Figure 6).

### Estimation of SNV mutation rates per base per cell division

Mutation rates per haploid base per cell division were calculated as described below. *De novo* indels and *de novo* SNVs on the X chromosome (in humans) were removed before analysis. MNVs (DNM sites greater or less than 5bp apart) were counted as a single event. The number of bases at which mutations could have been called was determined on an individual-specific basis as described previously^9^ taking into account both hard filters (e.g. repetitive sequences) and sequence depth in each individual. The average number of bases per genome at which mutations could have been detected was calculated to be 2,222,635,788bp in mice and 2,394,138,713bp in humans. Below, we used the number of mutations per offspring adjusted for the partial contribution of VEE mutations to the germline, as described above.

#### Average mutation rates per cell division

Average mutation rates per base per cell division were calculated by dividing the average number of mutations per offspring by the number of bases in the genome that were interrogated, and the average of the total paternal and maternal cell divisions calculated to have occurred in the offspring^11^, and divided by two to obtain a haploid mutation rate. The 95% conference intervals were calculated assuming the numbers of mutations per offspring are Poisson distributed.

To calculate the average paternal mutation rate per base per cell division, the mean number of autosomal mutations per offspring was first scaled by the proportion of phased mutations that are of paternal origin, to give the mean number of paternal autosomal mutations per generation, which was then divided by the estimated number of paternal cell divisions per generation (62 in mice, 401 in humans)^11^, and the number of bases interrogated in the genome. 95% confidence intervals were derived assuming Poisson variance. The average maternal mutation rate per base per cell division average was calculated similarly, using 25 and 31 cell divisions per generation (mouse and human, respectively)^11^.

#### Mutation rate per base per cell division for Very Early Embryonic Mutations

Very early embryonic (VEE) mutations occur in the first cell divisions that contribute to the embryo (rather than to extra-embryonic tissues). Modelling (assuming symmetric contributions of daughter cells to the embryo) shows that our mutation calling pipeline on WGS data only had substantial power to detect VEE mutations occurring in the first cell division and therefore present in ∼25% of reads, and had very low power to detect VEE mutations in subsequent cell divisions (12.5% of reads and lower). Moreover, the distribution of the Variant Allele Fraction for VEE mutations is centred symmetrically around 0.25 (Extended Data Figure 6) as expected for mutations arising in the first cleavage cell division contributing to the embryo. These results suggest that the vast majority of VEE mutations we detected arose in a single cell division.

To estimate the VEE mutation rate per base per cell division, we divided the mean number of VEE mutations per offspring, by the number of bases interrogated in the genome and then divided this number by 2 (to obtain a haploid rate). To allow for over-dispersion in VEE mutation rates, we calculated the 95% confidence interval assuming quasi-Poisson distribution^33^ in the total number of VEE mutations.

#### Mutation rate per base per cell division pre-puberty in the male germline

The mean number of mutations occurring pre-puberty in the male germline were estimated by subtracting the number of mutations expected to have accumulated since puberty, due to the parental age effect, from the total number of mutations observed in offspring as follows:

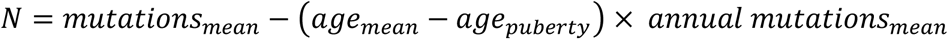

The age at puberty in humans and mice was assumed to be 15 years and 1 month respectively. The increase in number of mutations per additional year of parental age in humans was taken from the linear model described by Kong et al^2^. The increase in number of mutations per additional year of parental age in mice was extrapolated from the mean difference (6) in numbers of prezygotic mutations between first and last sequenced litters (separated by 33 weeks). The mean number of pre-pubertal mutations per offspring were then divided by the number of bases interrogated in the genome and the expected number of pre-pubertal cell divisions. 95% confidence intervals were derived from the standard error around the slope of the linear model describing the effect of age on the number of DNMs fitted to the data (human data obtained from Kong et al^2^).

#### Mutation rate per base per cell division post-puberty in the male germline

Mutations accrue in an approximately linear manner with parental age. The post-puberty mutation rate per base per cell division in males was estimated by dividing estimates of the increase in number of mutations per additional year of parental age (see above) by the estimated number of additional SSC divisions per year (42 for mice, 23 for humans^11^), and then by the number of bases interrogated in the genome. Confidence intervals were derived from the standard error around the slope of the linear model describing the effect of paternal age on the number of DNMs fitted to the data (human data obtained from Kong et al^2^).

#### Assessing significance in differences between mutation rates

Where count data were available, significance between mutation rates per base per cell division was tested using Wilcoxon rank test, and Students t-test. Where only mean and confidence intervals were available, significance was tested using the t-test only. All tests were adjusted for multiple testing using Bonferroni correction.

### Calculating Parental age effect

We observed an average increase of 6 DNMs over the 33 weeks between earliest and latest mouse litters in the pedigrees the two largest pedigrees, which is approximately five-fold greater (p= 0.0003) than we would expect in humans over the same time period ^2,9,10,11^. The remaining four pedigrees have information only from one time point in between the earliest and latest litters (Extended Table 1). We therefore combined all six pedigrees and constructed a mixed effects-linear model with the pedigree as a random effect to account for differences between pedigrees.

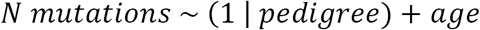

### Reconstruction of parental lineages

Parental lineages were reconstructed using the distribution of mutations shared between offspring, using an iterative algorithm derived from the following three expectations: This iterative procedure is based on three simple expectations: (i) mutations arising in different parents should co-occur in offspring at random, (ii) mutations present in the same diploid progenitor cell should co-occur in offspring more frequently than expected by chance, and (iii) mutations arising in the same parent but in different progenitor cells should be observed mutually exclusively in offspring (Methods).

In the first step, mutations belonging to the same parental lineage were identified iteratively using a pairwise test for all pairs of shared mutations, which calculated the binomial probability of ***n*** pups sharing ***m*** mutations where the frequencies of the mutations were ***p*** and ***q*** in the offspring. In each step of the iteration, the pair of mutations with the most significant p-value were considered to belong to the same parental lineage, as long as the parental origin of the two mutations was not discordant, and were merged into a single ‘pseudo-mutation’ assigned to all the offspring carrying either mutation. This procedure was then repeated iteratively, with each step involving merging of mutations belonging to the same parental lineage, either until a given p-value threshold is reached, or the pseudo mutations cannot be merged any further. Using a p-value threshold of 0.05, all mutations had completely collapsed into the clusters described. All but four of the seventy shared mutations could be assigned to either paternal or maternal lineages.

The accuracy of the lineage reconstruction algorithm was assessed using simulations. Firstly, for each pedigree, shared mutations were randomly re-assigned into the lineages defined by the reconstruction above. The pattern of mutation sharing was then assessed for biological concordance - each offspring can only belong to one paternal and one maternal lineage. The random reassignment of mutations was carried out 10,000 times for each pedigree, but none of these were biologically concordant (ie at least one offspring would have more than one paternal or maternal lineage). Secondly, for each pedigree, mutations were randomly clustered into lineages containing differing numbers of mutations (from 2-10 variant sites) and tested again for concordance as above, 10,000 times. In this way, 40,000 simulations across both pedigrees showed no other possible concordant lineage structures. Using this iterative clustering procedure, we assigned 67/70 shared mutations to a specific parent, and defined partial cellular genealogies for each parent. These primary lineages are distributed randomly with respect to litter timing, suggesting that their relative representation among gametes is stable over time and primarily reflects processes operating prior to PGC specification and/or during the early stages of PGC proliferation

## Supplementary Note 1

We validated 366 unique *de novo* mutations in CBGP4, CBGP7, GPCB9 and CBGP11 including 4 MNVs and 362 SNVs. We found 38/366 (13%) unique DNMs were EEs in the four pedigrees ranging from 1-13% of parental somatic cells. We detected 31/366 (8%) unique DNMs as peri-PGC (Paternal:Maternal 7:7 read pair phasing). Lastly we observed 205/386 observed DNMs as late-post PGC mutations. 92/386 observed DNMs (23%) were VEEs, also showed high variance of 1-11 DNMs per individual, more than expected under a Poission distribution (p=0.037), with all pedigrees combined (p=0.0019). They arose at the same rates in male and female offspring 4:4 and at similar rates on parental haplotypes 3:4 (Maternal:Paternal).

### Extended Data Tables and Figures

**Extended Data Table 1.** Mouse DNM counts. Table showing counts of DNMs in each strata for each individual. Offspring are labelled as a-g in litters ^*^_1 to ^*^_9. *De novo* indels were excluded from the analyses. The column “corrected SNVs” shows the number of autosomal SNVs corrected for the areas of the genome inaccessible to our WGS study, and for likely inheritance of VEEs.

| ID | Early Embryonic | peri-PGC | late-postPGC | VEE | total | Corrected total | sex | Age of parents at conception of offspring |
| --- | --- | --- | --- | --- | --- | --- | --- | --- |
| CBGP4_6a | 2 | 1 | 8 | 4 | 15 | 14.55 | M | 37 |
| CBGP4_6b | 2 | 2 | 9 | 6 | 19 | 16.74 | M | 37 |
| CBGP4_6c | 3 | 1 | 8 | 2 | 14 | 14.39 | M | 37 |
| CBGP4_6d | 2 | 1 | 13 | 4 | 20 | **19.05** | M | 37 |
| CBGP4_6e | 5 | 2 | 12 | 3 | 22 | **22.39** | F | 37 |
| CBGP7_2a | 3 | 3 | 12 | 3 | 21 | **21.12** | M | 12 |
| CBGP7_2b | 4 | 3 | 10 | 4 | 21 | **20.63** | M | 12 |
| CBGP7_2c | 5 | 2 | 7 | 7 | 21 | **18.26** | M | 12 |
| CBGP7_2d | 5 | 2 | 12 | 6 | 25 | **23.69** | M | 12 |
| CBGP7_2e | 2 | 2 | 6 | 11 | 21 | **17.56** | F | 12 |
| CBGP8_1a | 1 | 3 | 6 | 5 | 15 | **14.03** | M | 8 |
| CBGP8_1b | 0 | 5 | 6 | 7 | 18 | **16.47** | M | 8 |
| CBGP8_1c | 2 | 5 | 8 | 4 | 19 | **18.28** | M | 8 |
| CBGP8_1g | 1 | 6 | 4 | 5 | 16 | **14.50** | F | 8 |
| CBGP8_1h | 1 | 1 | 10 | 2 | 14 | **14.17** | F | 8 |
| CBGP8_8a | 1 | 3 | 13 | 1 | 18 | **19.16** | M | 40 |
| CBGP8_8b | 2 | 3 | 8 | 5 | 18 | **17.15** | M | 40 |
| CBGP8_8c | 0 | 2 | 12 | 5 | 19 | **18.01** | M | 40 |
| CBGP8_8d | 3 | 5 | 19 | 9 | 36 | **33.53** | F | 40 |
| CBGP8_8f | 4 | 5 | 11 | 3 | 23 | **23.88** | F | 40 |
| GPCB11_1a | 1 | 3 | 11 | 3 | 18 | 17.77 | M | 8 |
| GPCB11_1b | 0 | 2 | 7 | 7 | 16 | 13.45 | M | 8 |
| GPCB11_1d | 2 | 3 | 2 | 1 | 8 | 8.68 | M | 8 |
| GPCB11_1f | 4 | 1 | 8 | 5 | 18 | 15.80 | F | 8 |
| GPCB11_1g | 3 | 1 | 15 | 1 | 20 | 21.31 | F | 8 |
| GPCB2_1a | 1 | 0 | 10 | 13 | 24 | **19.26** | M | 9 |
| GPCB2_1b | 1 | 4 | 5 | 4 | 14 | **12.75** | M | 9 |
| GPCB2_1c | 1 | 1 | 15 | 8 | 25 | **23.53** | M | 9 |
| GPCB2_1d | 1 | 9 | 11 | 3 | 24 | **24.38** | M | 9 |
| GPCB2_1e | 1 | 6 | 5 | 1 | 13 | **13.88** | M | 9 |
| GPCB2_9a | 2 | 8 | 12 | 4 | 26 | **24.95** | M | 41 |
| GPCB2_9b | 1 | 2 | 9 | 6 | 18 | **16.02** | M | 41 |
| GPCB2_9c | 0 | 5 | 16 | 7 | 28 | **26.86** | M | 41 |
| GPCB2_9e | 2 | 3 | 22 | 0 | 27 | **29.84** | M | 41 |
| GPCB2_9f | 2 | 3 | 15 | 10 | 30 | **26.58** | F | 41 |
| GPCB9_8a | 0 | 2 | 20 | 11 | 33 | 28.03 | M | 40 |
| GPCB9_8c | 1 | 1 | 12 | 3 | 17 | 16.81 | F | 40 |
| GPCB9_8d | 2 | 0 | 8 | 3 | 13 | 12.39 | F | 40 |
| GPCB9_8f | 2 | 3 | 17 | 5 | 27 | 28.40 | F | 40 |
| GPCB9_8g | 0 | 3 | 11 | 3 | 17 | 16.34 | F | 40 |

**Extended Data Table 2.** Sharing of DNMs between WGS offspring in the two largest mouse pedigrees. The probability of a DNM being present in more than one offspring in the same pedigree was calculated as the number of instances of a mutation being shared by two offspring in the same pedigree divided by the number of pairwise comparisons between offspring in each pedigree. (for further details on methodology and human data see Rabhari et al^9^)

| Pedigree | instances of sib-sib sharing DNMs | total comparisons | Number of shared DNMs per offspring |
| --- | --- | --- | --- |
| CBGP8 | 90 | 1800 | 5 |
| GPCB2 | 32 | 2151 | 1.49 |
| total | 122 | 3951 | 3.09 |

**Extended Data Table 3.** Early Embryonic DNMs. Early Embryonic DNMs from six mouse pedigrees, annotated with parent of origin and levels of somatic and germline mosaicism observed. Germline mosaic frequency was calculated as number of offspring carrying mutation/total number of offspring assayed.

| Pedigree | Chr | Pos | Ref | Alt | Mosaic Parent | mean % of reads with alternative allele in parental tissues | Number of offspring with mutation | Number of offspring sampled | LINEAGE | germline mosaic frequency |
| --- | --- | --- | --- | --- | --- | --- | --- | --- | --- | --- |
| CBGP4 | 2 | 41430847 | C | T | Father | 2.342 | 6 | 41 | NA | 0.146 |
| CBGP4 | 2 | 53187077 | T | C | Father | 10.371 | 7 | 41 | NA | 0.171 |
| CBGP4 | 3 | 33383613 | A | T | Mother | 3.366 | 1 | 41 | NA | 0.024 |
| CBGP4 | 6 | 34892512 | G | A | Father | 5.496 | 2 | 41 | NA | 0.049 |
| CBGP4 | 12 | 64770094 | T | G | Mother | 4.548 | 7 | 41 | NA | 0.171 |
| CBGP4 | 12 | 96555454 | T | G | Father | 12.668 | 2 | 41 | NA | 0.049 |
| CBGP4 | 14 | 97813384 | G | A | Father | 1.193 | 1 | 41 | NA | 0.024 |
| CBGP4 | 15 | 12730537 | C | A | Father | 10.601 | 9 | 41 | NA | 0.22 |
| CBGP4 | 15 | 29094924 | A | T | Father | 9.975 | 2 | 41 | NA | 0.049 |
| CBGP4 | 18 | 59431259 | G | T | Father | 1.644 | 1 | 41 | NA | 0.024 |
| CBGP4 | 18 | 63653717 | C | T | Father | 3.094 | 1 | 41 | NA | 0.024 |
| CBGP4 | 18 | 77628350 | C | G | Father | 0.919 | 1 | 41 | NA | 0.024 |
| CBGP7 | 1 | 3219586 | G | A | Father | 11.875 | 14 | 46 | NA | 0.341 |
| CBGP7 | 1 | 109973541 | A | T | Father | 5.392 | 9 | 46 | NA | 0.22 |
| CBGP7 | 2 | 11064217 | G | C | Father | 5.563 | 6 | 46 | NA | 0.146 |
| CBGP7 | 2 | 18380082 | A | G | Father | 12.91 | 12 | 46 | NA | 0.293 |
| CBGP7 | 3 | 43034668 | T | G | Father | 2.295 | 2 | 46 | NA | 0.049 |
| CBGP7 | 4 | 47685275 | T | C | Mother | 1.19 | 12 | 46 | NA | 0.293 |
| CBGP7 | 5 | 123117988 | C | T | Father | 2.528 | 7 | 46 | NA | 0.171 |
| CBGP7 | 8 | 118892306 | G | T | Mother | 1.297 | 10 | 46 | NA | 0.244 |
| CBGP7 | 9 | 82120658 | C | T | Mother | 2.638 | 11 | 46 | NA | 0.268 |
| CBGP7 | 17 | 8698893 | G | A | Father | 12.678 | 13 | 46 | NA | 0.317 |
| CBGP7 | 17 | 48347925 | T | G | Father | 2.023 | 4 | 46 | NA | 0.098 |
| CBGP7 | 18 | 82381825 | A | G | Father | 7.455 | 4 | 46 | NA | 0.098 |
| CBGP7 | 3 | 70223928 | G | A | Father | 2.213 | 1 | 46 | private | 0.024 |
| CBGP8 | 2 | 10556273 | G | A | Father | 2.612 | 10 | 57 | P1 | 0.244 |
| CBGP8 | 2 | 92472946 | A | T | Father | 2.122 | 10 | 57 | P1 | 0.244 |
| CBGP8 | 4 | 28191752 | T | C | Father | 4.05 | 4 | 57 | P3 | 0.098 |
| CBGP8 | 10 | 89180326 | A | T | Father | 9.508 | 10 | 57 | P2 | 0.244 |
| CBGP8 | 13 | 61390887 | T | C | Father | 1.458 | 2 | 57 | P3 | 0.049 |
| CBGP8 | 15 | 60633937 | A | C | Father | 2.913 | 7 | 57 | P1 | 0.171 |
| CBGP8 | 17 | 64534541 | G | C | Father | 2.342 | 12 | 57 | P1 | 0.293 |
| CBGP8 | 17 | 89395091 | G | C | Father | 7.695 | 1 | 57 | private | 0.024 |
| GPCB11 | 1 | 138426118 | T | C | Father | 10.834 | 10 | 47 | NA | 0.244 |
| GPCB11 | 2 | 142507389 | G | A | Father | 10.061 | 10 | 47 | NA | 0.244 |
| GPCB11 | 2 | 146721803 | G | A | Father | 9.818 | 9 | 47 | NA | 0.22 |
| GPCB11 | 3 | 83756086 | C | T | Mother | 2.684 | 4 | 47 | NA | 0.098 |
| GPCB11 | 8 | 110980784 | G | A | Mother | 2.581 | 4 | 47 | NA | 0.098 |
| GPCB11 | 12 | 53018942 | G | T | Mother | 1.296 | 7 | 47 | NA | 0.171 |
| GPCB11 | 15 | 22732768 | C | T | Mother | 1.926 | 4 | 47 | NA | 0.098 |
| GPCB11 | 18 | 32504139 | G | A | Father | 3.344 | 6 | 47 | NA | 0.146 |
| GPCB2 | 1 | 195105306 | C | T | Father | 1.905 | 8 | 77 | P2 | 0.195 |
| GPCB2 | 2 | 123112871 | G | A | Father | 4.795 | 9 | 77 | P3 | 0.22 |
| GPCB2 | 3 | 27601982 | A | C | Father | 6.166 | 6 | 77 | P4 | 0.146 |
| GPCB2 | 4 | 88898019 | T | A | Mother | 4.093 | 10 | 77 | M1 | 0.244 |
| GPCB2 | 6 | 76705056 | A | G | Father | 0.938 | 7 | 77 | P4 | 0.171 |
| GPCB2 | 11 | 56513226 | A | T | Father | 2.512 | 12 | 77 | P3 | 0.293 |
| GPCB2 | 11 | 67581517 | G | A | Father | 9.218 | 9 | 77 | P2 | 0.22 |
| GPCB2 | 12 | 14764225 | C | T | Father | 7.395 | 9 | 77 | P2 | 0.22 |
| GPCB2 | 18 | 68245753 | G | T | Father | 0.824 | 13 | 77 | P1 | 0.317 |
| GPCB9 | 14 | 14762737 | T | G | Mother | 3.739 | 2 | 77 | NA | 0.049 |
| GPCB9 | 14 | 78127137 | A | T | Mother | 3.846 | 2 | 77 | NA | 0.049 |
| GPCB9 | 15 | 22224009 | A | G | Mother | 1.363 | 1 | 77 | NA | 0.024 |
| GPCB9 | 18 | 88094408 | C | A | Mother | 7.681 | 1 | 77 | NA | 0.024 |
| GPCB9 | 19 | 34151974 | G | A | Father | 10.926 | 3 | 77 | NA | 0.073 |

**Extended Data Table 4:** DNMs with potentially functional consequences as given by ANNOVAR ^29^ are listed.

| IIDs | chr | position | reference | alternative | consequence | gene | strata |
| --- | --- | --- | --- | --- | --- | --- | --- |
| CBGP8_1k, CBGP8_2a, CBGP8_2i,<br>CBGP8_5a, CBGP8_5b, CBGP8_5c,<br>CBGP8_7a, CBGP8_7c, CBGP8_8b,<br>CBGP8_8d | 2 | 10556273 | G | A | synonymous SNV | Sfmbt2 | paternal early<br>embryonic |
| CBGP8_8f | 2 | 28556769 | A | T | nonsynonymous<br>SNV | Cel | late post-PGC<br>specification |
| CBGP8_1a, CBGP8_3a, CBGP8_4a,<br>CBGP8_4h, CBGP8_5d | 2 | 30084760 | C | T | nonsynonymous<br>SNV | Pkn3 | paternal peri-PGC specification |
| CBGP8_8a | 7 | 104975169 | T | C | nonsynonymous<br>SNV | Olfr671 | late post-PGC<br>specification |
| CBGP8_1c | 8 | 83722794 | G | A | splicing | Ddx39 | late post-PGC<br>specification |
| CBGP8_1g | 9 | 123480538 | C | T | nonsynonymous<br>SNV | Limd1 | late post-PGC<br>specification |
| GPCB2_1d | 11 | 50873775 | G | A | stopgain | Zfp454 | late post-PGC<br>specification |
| CBGP8_1h | 13 | 100154877 | C | T | synonymous SNV | Naip2 | late post-PGC<br>specification |
| CBGP8_1h | 13 | 100154880 | G | A | synonymous SNV | Naip2 | late post-PGC<br>specification |
| CBGP8_1h | 13 | 100154911 | T | A | nonsynonymous SNV | Naip2 | late post-PGC specification |
| CBGP8_1h | 13 | 100154951 | C | T | nonsynonymous SNV | Naip2 | late post-PGC specification |
| CBGP8_1a | 19 | 44935143 | G | A | nonsynonymous SNV | Fam178a | late post-PGC specification |
| CBGP7_2e | 2 | 76889786 | T | C | nonsynonymous SNV | Ttn | VEE |
| CBGP4_6e | 3 | 146944022 | G | A | synonymous SNV | Ttll7 | VEE |
| CBGP4_6c,CBGP4_6e | 6 | 34892512 | G | A | nonsynonymous SNV | Wdr91 | paternal early embryonic |
| GPCB9_8g | 7 | 12417167 | T | C | nonsynonymous SNV | Zfp551 | late post-PGC specification |
| GPCB11_1d,GPCB11_4c | 7 | 27465841 | C | T | synonymous SNV | Blvrb | peri-PGC specification |
| GPCB9_8f | 9 | 106224354 | T | A | nonsynonymous SNV | Tlr9 | late post-PGC specification |
| GPCB9_8c | 10 | 5075830 | C | T | nonsynonymous SNV | Syne1 | late post-PGC specification |
| CBGP7_2a,CBGP7_2b,CBGP7_6b,CBGP7_8e | 10 | 81261530 | C | T | nonsynonymous SNV | Matk | peri-PGC specification |
| CBGP7_2a,CBGP7_2b | 11 | 115802091 | G | A | nonsynonymous SNV | Caskin2 | peri-PGC specification |
| CBGP7_2a | 13 | 105250190 | C | T | synonymous SNV | Rnf180 | late post-PGC specification |
| CBGP4_6a | 17 | 24600010 | A | C | nonsynonymous SNV | Tsc2 | late post-PGC specification |
| CBGP7_2e | 17 | 25492005 | G | A | synonymous SNV | Sstr5 | VEE |
| CBGP7_2e | 18 | 64561658 | A | C | nonsynonymous SNV | Atp8b1 | late post-PGC specification |
| CBGP7_2b | 19 | 7281156 | G | T | synonymous SNV | Mark2 | late post-PGC specification |

**Extended Data Table 5:** Quantification of VEE mutations to the germline. All 40 putative parental VEE mutations that we attempted to validate are listed. Sites classed as “FAIL” either failed to generate informative data, or were not a VEE mutation.

| chr | pos | ref | alt | ped | Parent of origin | Outcome | Number of Offspring Assayed | Number of Offspring with mutation | Mosaicism | Average variant allele fraction (Offspring) | Variant allele fraction (Parents) |
| --- | --- | --- | --- | --- | --- | --- | --- | --- | --- | --- | --- |
| 5 | 40039235 | C | G | CBGP4 | Father | FAIL | NA | NA | NA | NA | NA |
| 9 | 9651062 | G | T | GPCB11 | Father | FAIL | NA | NA | NA | NA | NA |
| 9 | 51859263 | G | A | CBGP7 | Mother | FAIL | NA | NA | NA | NA | NA |
| 6 | 47509685 | A | T | GPCB11 | Father | FAIL | 47 | 0 | 0 | 0 | 0.07 |
| 1 | 112055045 | C | A | CBGP4 | Mother | FAIL | 40 | 2 | 0.05 | 0.71 | 0.16 |
| 2 | 58446583 | C | T | GPCB9 | Father | FAIL | 54 | 18 | 0.33 | 0.98 | 0.28 |
| 7 | 79891839 | T | C | GPCB9 | Mother | FAIL | 55 | 3 | 0.05 | 0.52 | 0.11 |
| 7 | 95769188 | C | A | GPCB9 | Mother | FAIL | 54 | 20 | 0.37 | 0.5 | 0.16 |
| 12 | 83079835 | A | C | GPCB11 | Mother | FAIL | 47 | 14 | 0.3 | 0.5 | 0.06 |
| 13 | 44372937 | T | G | GPCB11 | Mother | FAIL | 47 | 7 | 0.15 | 0.5 | 0 |
| 13 | 64463785 | G | A | CBGP7 | Father | FAIL | 45 | 4 | 0.09 | 0.49 | 0.09 |
| 17 | 59986828 | C | T | CBGP7 | Father | FAIL | 46 | 5 | 0.11 | 0.5 | 0.07 |
| 1 | 140558754 | G | A | CBGP7 | Father | PASS | 46 | 11 | 0.24 | 0.5 | 0.21 |
| 2 | 38136612 | T | A | CBGP7 | Father | PASS | 46 | 4 | 0.09 | 0.5 | 0.15 |
| 2 | 75299374 | T | A | CBGP4 | Father | PASS | 40 | 1 | 0.03 | 0.5 | 0.15 |
| 2 | 89393187 | T | A | CBGP7 | Mother | PASS | 46 | 13 | 0.28 | 0.5 | 0.13 |
| 3 | 59107159 | A | G | GPCB9 | Mother | PASS | 55 | 11 | 0.2 | 0.5 | 0.21 |
| 4 | 29438300 | T | C | GPCB11 | Father | PASS | 47 | 7 | 0.15 | 0.5 | 0.09 |
| 4 | 32934671 | C | T | CBGP7 | Father | PASS | 41 | 21 | 0.51 | 0.5 | 0.22 |
| 4 | 52046695 | G | C | GPCB11 | Mother | PASS | 47 | 1 | 0.02 | 0.51 | 0.07 |
| 5 | 146886001 | A | T | GPCB9 | Mother | PASS | 55 | 9 | 0.16 | 0.49 | 0.19 |
| 6 | 26640451 | A | G | CBGP4 | Father | PASS | 40 | 5 | 0.13 | 0.49 | 0.24 |
| 6 | 26971358 | C | G | GPCB11 | Father | PASS | 47 | 4 | 0.09 | 0.5 | 0.23 |
| 6 | 28151992 | A | G | GPCB9 | Mother | PASS | 55 | 9 | 0.16 | 0.44 | 0.2 |
| 6 | 149428040 | T | C | GPCB9 | Mother | PASS | 55 | 18 | 0.33 | 0.47 | 0.3 |
| 7 | 119861636 | C | T | GPCB9 | Mother | PASS | 54 | 16 | 0.3 | 0.51 | 0.31 |
| 8 | 59958418 | G | A | GPCB9 | Mother | PASS | 54 | 14 | 0.26 | 0.5 | 0.31 |
| 8 | 78366703 | G | C | CBGP4 | Mother | PASS | 40 | 4 | 0.1 | 0.5 | 0.16 |
| 8 | 87011056 | C | T | GPCB11 | Mother | PASS | 47 | 0 | 0 | 0 | 0.08 |
| 8 | 127818809 | A | G | GPCB9 | Mother | PASS | 54 | 7 | 0.13 | 0.49 | 0.21 |
| 10 | 50837657 | C | T | CBGP7 | Mother | PASS | 46 | 0 | 0 | 0 | 0.15 |
| 12 | 29010841 | G | T | CBGP4 | Father | PASS | 40 | 7 | 0.18 | 0.51 | 0.08 |
| 12 | 48054417 | C | T | GPCB9 | Father | PASS | 54 | 20 | 0.37 | 0.51 | 0.28 |
| 12 | 67115109 | A | G | CBGP4 | Mother | PASS | 50 | 4 | 0.08 | 0.48 | 0.15 |
| 14 | 122391093 | T | C | GPCB9 | Father | PASS | 55 | 0 | 0 | 0 | 0.09 |
| 6 | 72332353 | G | T | GPCB11 | Father | PASS | 46 | 0 | 0 | 0 | 0.04 |
| 9 | 80645537 | G | A | GPCB9 | Mother | PASS | 54 | 0 | 0 | 0 | 0.02 |
| 1 | 88802774 | C | G | GPCB9 | Father | PASS | 52 | 24 | 0.46 | 0.5 | 0.42 |
| 16 | 15153420 | G | T | GPCB9 | Father | FAIL | NA | NA | NA | NA | NA |
| 18 | 51620274 | A | G | GPCB9 | Mother | FAIL | NA | NA | NA | NA | NA |

**Extended Data Table 6: Validated DNMs observed in six mouse pedigrees.** All DNMs are listed, with columns in order of chromosome, position, type, reference allele, variant allele, which offspring they were called in (the suffix ‘T’ on the offspring ID, e.g. CBGP8_1aT, indicates that tail DNA rather than spleen DNA was assayed), the number of individuals the site is shared with, the temporal strata to which it was assigned, which lineage it belongs to, and finally read-pair haplotyping results. CBGP7_2a has information from only one tissue (tail) due to a QC problems with DNA from the speen.

### Extended Data Figures

**Extended Data Figure 1:**
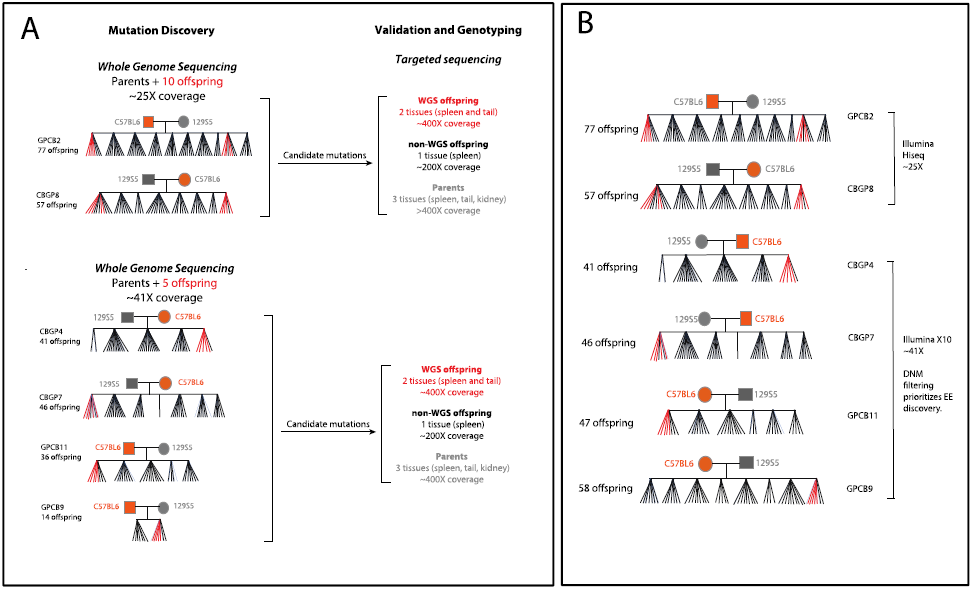
A) Mouse pedigree sequencing and genotyping strategy. Six reciprocal crosses were established and successively mated, generating six pedigrees comprising between 41 and 77 offspring (Extended Data Figure 1). The pedigrees shown here only include the sequenced and genotyped individuals. Three tissues (spleen, kidney and tail) were collected from all mice. Whole genome sequencing (WGS) was performed on spleen-derived DNA for five or ten pups within each pedigree (shown in red). Candidate DNMs were identified and validated using deeper targeted sequencing of 1-3 tissues per individual across the offspring shown. **B) Complete mouse pedigrees**. The naming conventions and differences between data collection and analysis is shown. Not all offspring from each pedigree were genotyped.

The naming conventions and the differences between data collection and analysis is shown. The offspring in red were whole genome sequenced. Not all offspring from each pedigree were genotyped.

**Extended Data Figure 2:**
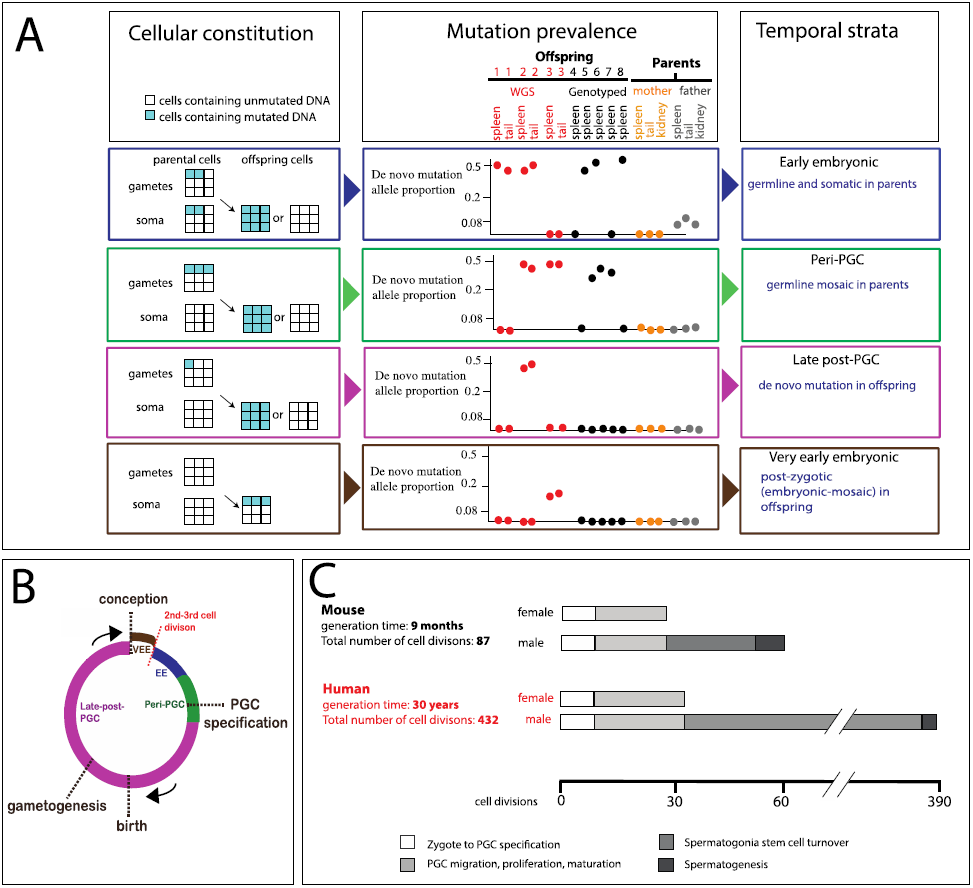
Temporal strata of observed mutations. **A.** (Left) Four classes of mutations distinguished by the proportion of cells carrying the mutation within parental and offspring tissues. Multiple tissues were used to verify consistency of VAF. (Middle) Illustrative examples of variant allele fraction observed among the sequenced samples for each class. (Right) Temporal strata to which these mutations are assigned. **B** Four temporal strata mapped onto the generational cycle of the germline: EE, peri-PGC and late post-PGC mutations were detected in parents, VEE mutations were detected in offspring. **C** Cell divisions occurring in the different stages of the mice and human germlines during an average generation^11^.

**Extended Data Figure 3:**
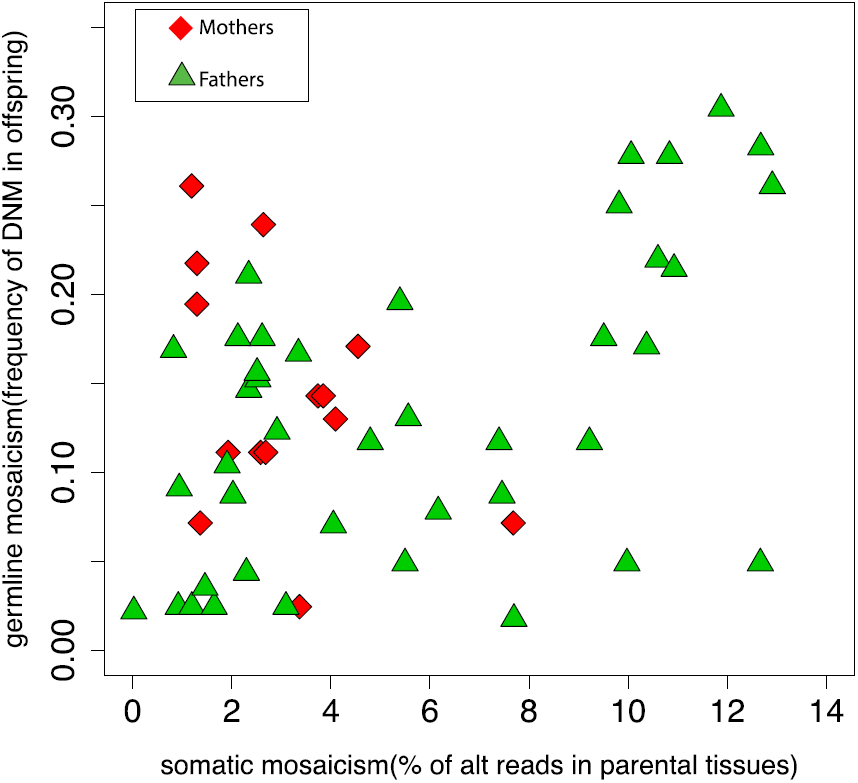
Levels of Somatic and Germline mosaicism observed at 55 EE mutations in six mouse pedigrees.

The y axis shows the germline mosaicism (number of DNMs in offspring/total number of offspring genotyped). The x axis shows the % of alternate reads observed in parental tissues.

**Extended Data Figure 4:**
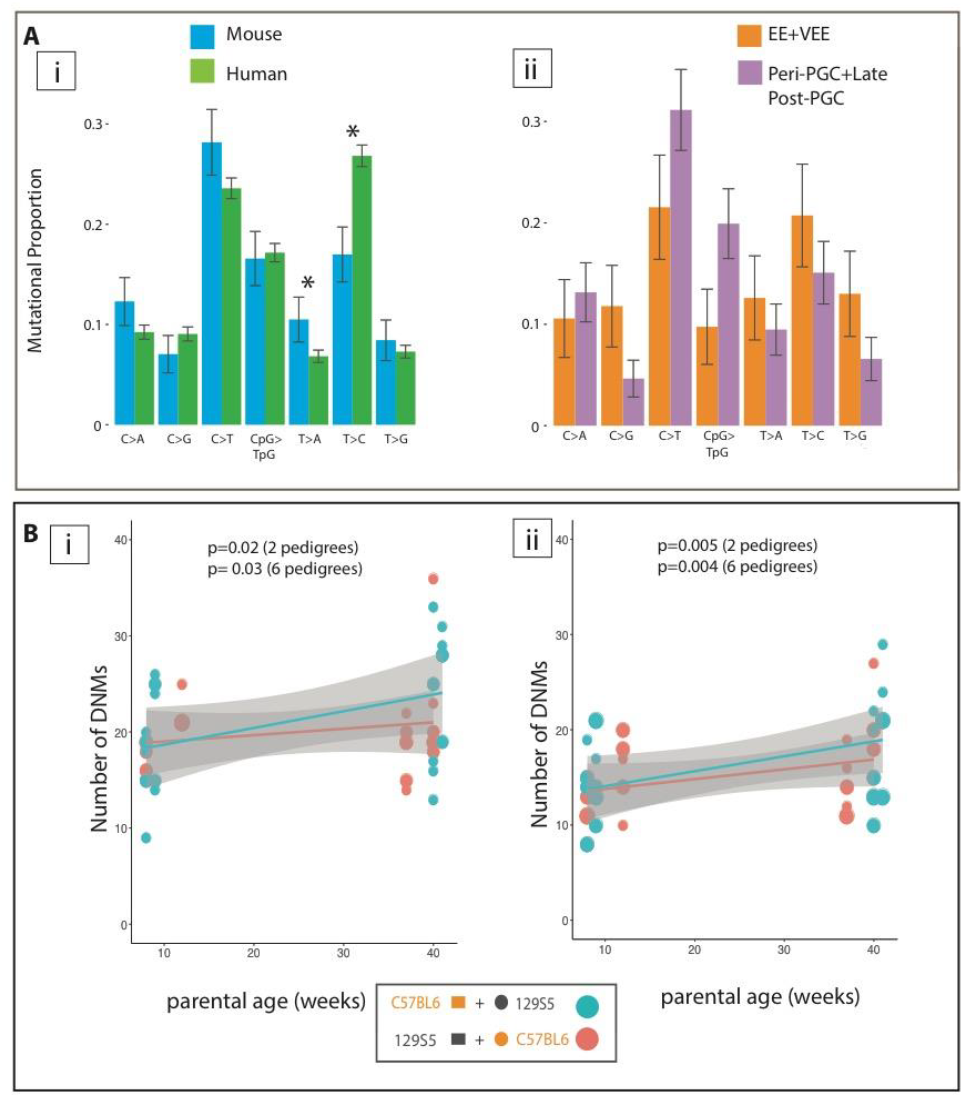
Mutational spectra and paternal age effect of DNMs in mice. A (i) Comparison of mutational spectra in mice and humans using a catalogue of compiled DNMs in humans^9^. Error bars are 95% confidence intervals. Stars show significance (p<0.05) after correction for multiple testing. (ii) Comparison of mutational spectra in VEE + EE mutations compared to Peri-PGC + late post-PGC mutations in mice. B) Effect of parental age on number of DNMs in individuals before (i) after (ii) removal of VEE mutations arising in offspring.

The size of the plotting symbol indicates the number of individuals sharing the same number of mutations. P values indicate the significance of the regression coefficients.

**Extended Data Figure 5:**
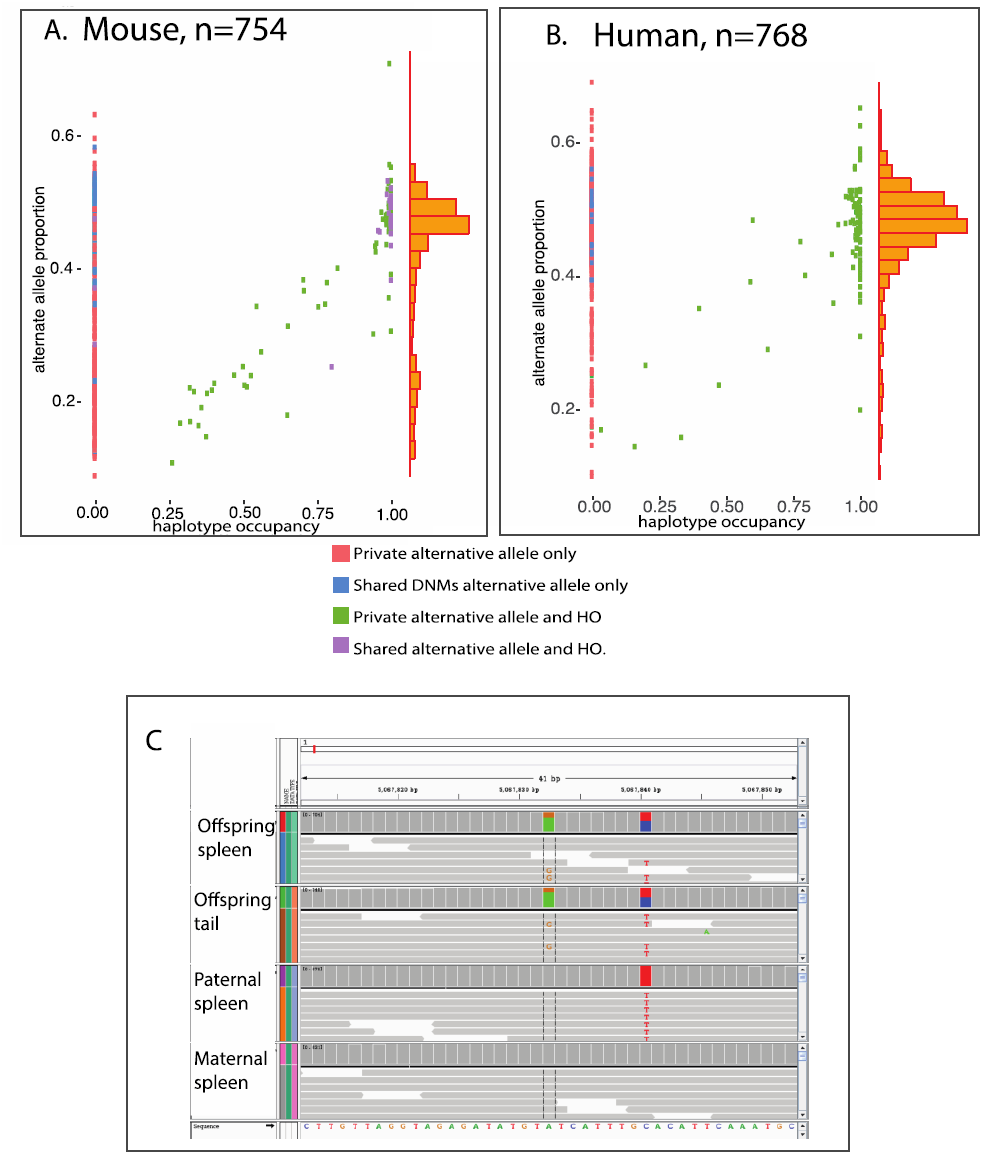
Haploptype Occupancy. A) Haplotype occupancy of DNMs in mice (haplotypes defined by adjacent heterozygous variants, Methods) plotted against the Variant Allele Fraction of the validated DNM. The right-hand histogram shows the distribution of VAF for DNMs for which haplotype occupancy could be determined. B) Haplotype occupancy of DNMs in humans, C) An example of incomplete haplotype occupancy defined as a DNM (in this case, A->G), which is not present on every instance of the haplotype (defined by the C->T variant) on which it arose.

**Extended Data Figure 6:**
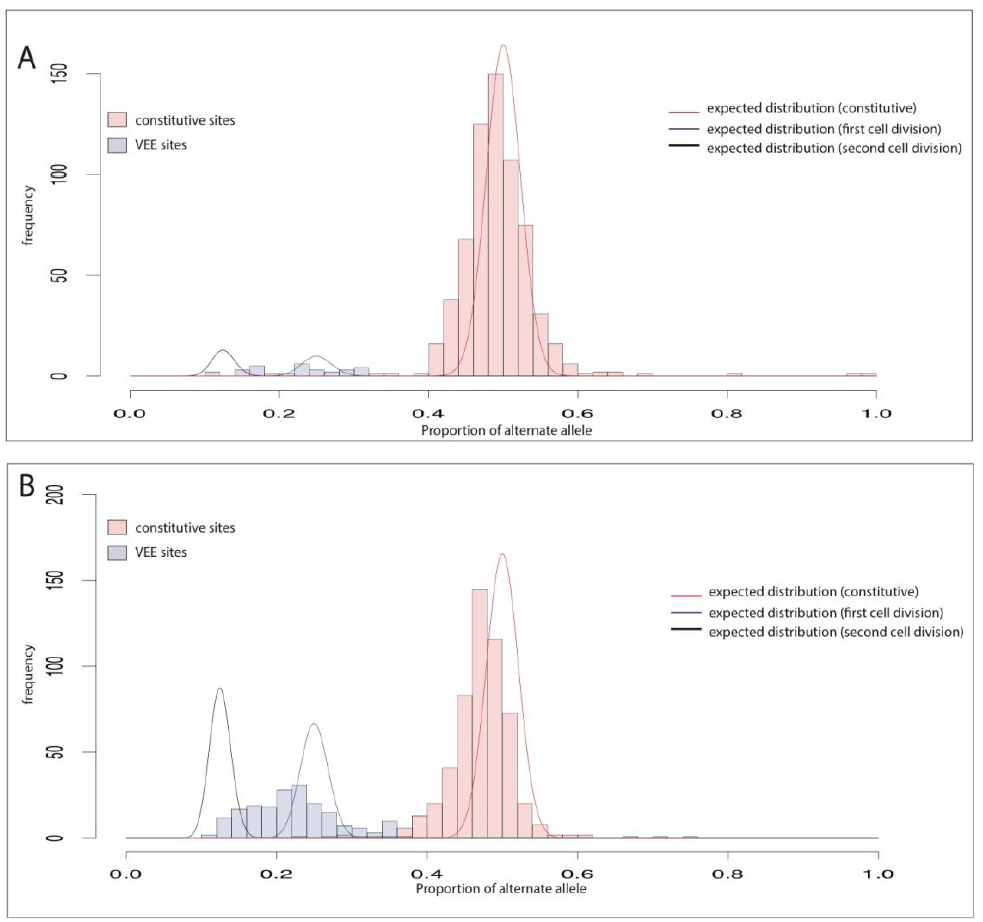
Variant Allele Fractions in Deep Sequenced Data. Histograms of the variant allele fraction in validated DNMs in the deep sequence data in humans (A) and mice (B). Constitutive sites are in red and have a VAF centred around 50% of reads (100% of cells). Very early embryonic (VEE) are shown in blue, are found in around 25% of reads (50% of cells). Red, blue and black lines show the expected distribution of VAF given a binomial distribution of reads centred around constitutive, first division and second division mutations (assuming symmetric contributions to the embryo), in our validation sequence data for all pedigrees.

**Extended Data Figure E7:**
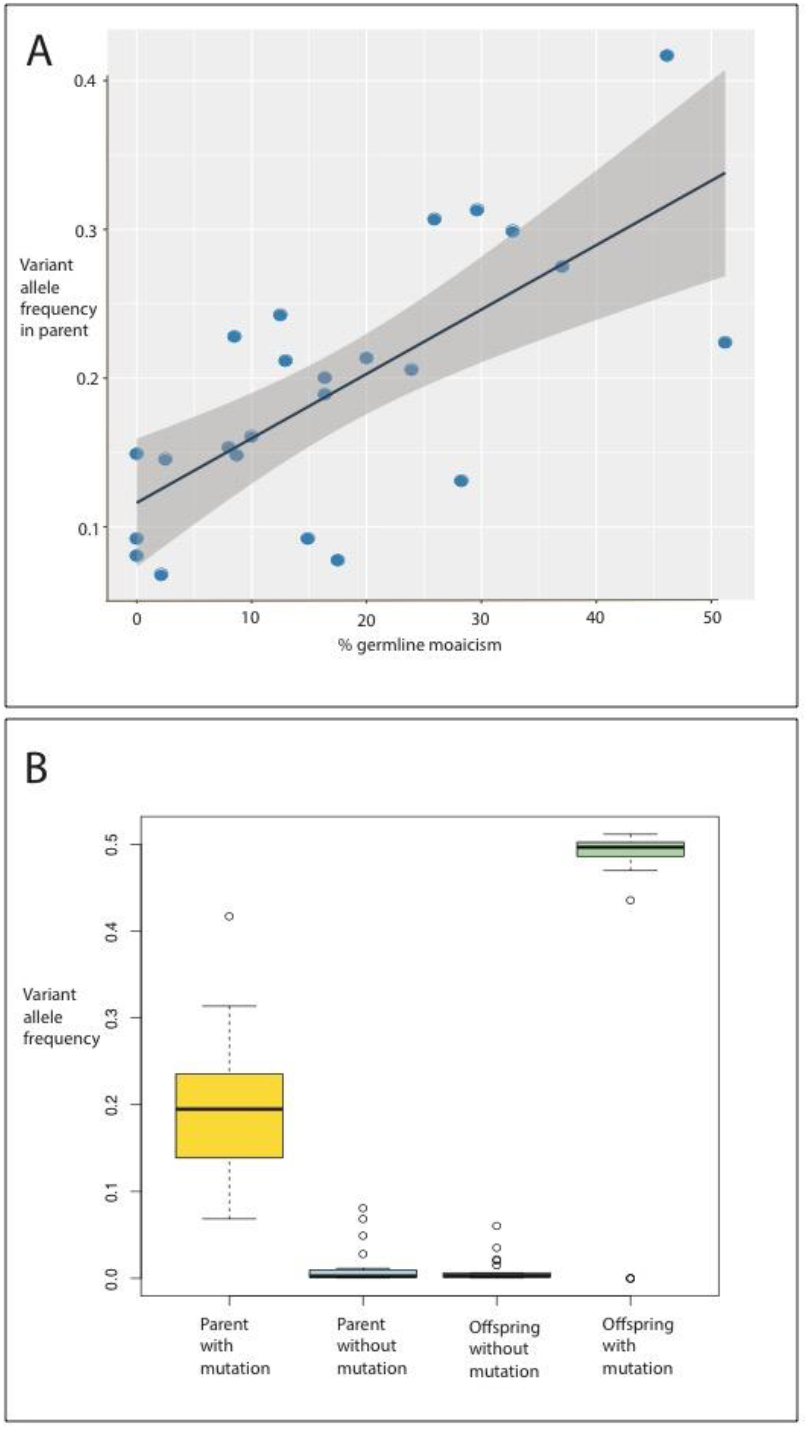
Germline mosaicism of VEE mutations observed in the parents. A) The frequency of the variant allele in the parent is plotted against the % of offspring assayed carrying the variant allele as an inherited constitutive variant. B) Boxplots of the frequency of the variant allele in the parents and offspring, demonstrating the distinction between the variant allele frequency observed as embryonic mutations in the parents and as constitutive variants in the offspring.

**Extended Data Figure 8:**
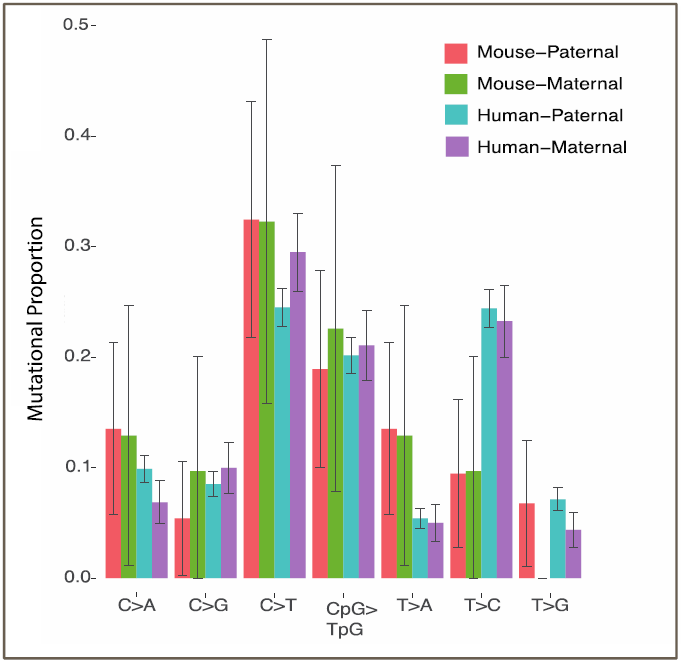
Low resolution mutation spectra in maternal and paternally derived DNMs in mouse and human^10^ data. Error bars show the 95% confidence intervals.

### Supplementary Note 1

We validated 366 unique *de novo* mutations in the CBGP4, CBGP7, GPCB9 and GPCB11 including 4 MNVs and 362 SNVs. We found 38/366 (13%) unique DNMs were EEs in the four pedigrees ranging from 1-13% of parental somatic cells. We detected 31/366 (8%) unique DNMs as peri-PGC (Paternal:Maternal 7:7 read pair phasing). Lastly we observed 205/386 observed DNMs as late-post PGC mutations. 92/386 observed DNMs (23%) were VEEs, also showed high variance of 1-11 DNMs per individual, more than expected under a Poission distribution (p=0.037), all pedigrees combined (p=0.0019). They arose at the same rates in male and female offspring 4:4 and at similar rates on parental haplotypes 3:4 (Maternal:Paternal).

## Acknowledgements

We are very grateful for the expert assistance of James Bussell and the Sanger Institute Mouse Facility for mouse breeding. We thank Mike Stratton, Inigo Martincorena, Art Wuster, Saeed Al-Turki, Jeremy McRae, Ludmil Alexandrov, Aylwyn Scally, Kirstie Lawson, Ian Adams and Robin Lovell-Badge for thoughtful discussions and sharing of scripts. This work was supported by the Wellcome Trust [grant number WT098051].

## Author contributions

The study was devised by SJL and MEH. Sample preparation was performed by SJL. Analyses were performed by SJL, RR, JK, TK and MEH. The manuscript was written by SJL, JK and MEH. The project was supervised by MEH.

## Accessions

The whole genome sequences generated as part of this study were deposited in the European Nucleotide Archive (ENA), under study PRJEB1407 and PRJEB14877.

